# Post-mortem infant-directed behaviours in wild Guinea baboons

**DOI:** 10.64898/2026.02.18.706182

**Authors:** Anaïs Avilés de Diego, Federica Dal Pesco, Julia Fischer

**Author notes:** Author responsible for correspondence: Anaïs Avilés de Diego, Address: Cognitive Ethology Laboratory, German Primate Centre, GmbH, Kellnerweg 4, 37077 Göttingen, Germany.

## Abstract

The growing field of comparative thanatology aims to shed light on how and why the understanding of death evolved. Observations across different nonhuman primate species have reported care-taking behaviour of dead infants, but also cannibalism. Various hypotheses have been put forward to explain these infant-directed behaviours, ranging from responses to infantile cues to an understanding of death. To aid comparative analyses and test some of these hypotheses, we report behaviours directed at dead infants in a wild population of Guinea baboons (*Papio papio*) living in the Niokolo-Koba National Park, Senegal. During 12 years of field observations (2014-2025), 67 infants died before reaching 1 year of age. In 4 cases, we could not establish when the infants had died because the field station was closed due to the COVID-19 pandemic. In 22 of the remaining 63 cases, mothers, but occasionally also other group members, carried, protected, groomed, and dragged dead infants. In 6 cases, cannibalism occurred. None of the mothers expressed any signs of emotional distress in response to infant death. We suggest that a concept of death in Guinea need not be invoked to explain the observed behaviours. Instead, selection appears to have favoured post-mortem caretaking behaviours to avoid abandoning an infant that might temporarily be unresponsive. The lack of infant responses to maternal behaviour and the disintegration of the corpse may drive the transition from perceiving the infants as an object that evokes caretaking to one that resembles food, which ultimately facilitates occurrences of cannibalism.

## Introduction

Over recent decades, growing interest in animal cognition and sentience, together with an increasing number of field and captive observations of behaviours directed towards dead individuals (hereafter ‘post-mortem behaviours’), have triggered the question of whether animals possess a basic awareness of death and whether such awareness might have contributed to the evolution of human sophisticated understanding of death (Anderson, 2017). Comparative thanatology has been defined as the scientific study of death and dying across animal taxa, and the behavioural, psychological, physiological, and social responses that accompany the death of conspecifics (Anderson, 2016, 2017). Studying the effects of the loss of a relative or a social partner can elucidate the evolutionary trajectories underlying death-related responses, from ancestral mechanisms to more derived traits.

Post-mortem behaviours, more specifically care-taking behaviours and carrying of dead individuals, have been observed in a wide variety of primate taxa (e.g., chimpanzees, *Pan troglodytes*: Bersacola et al., 2025; chacma baboons, *Papio ursinus*: Carter et al., 2020; Sumatran orangutans, *Pongo abelii*: Dellatore et al., 2009; northern muriquis, *Brachyteles hypoxanthus*: Freire Filho et al., 2022; Indian langurs, *Presbytis entellus*: Jay, 1962; mountain gorillas, *Gorilla beringei beringei*: Warren & Williamson, 2004). In these studies, the most frequently observed behaviours are inspection, protection, grooming, and carrying directed by mothers towards their dead infants, although occasionally other conspecifics may also direct post-mortem behaviours towards the infants (Carter et al., 2020; Merz, 1978). Whereas some mothers abandon their deceased infants soon after they die, others continue interacting with them for extended periods of time (>10 days; Fashing et al., 2011).

The most prominent post-mortem behaviour in primates is carrying (reviewed in Fernández-Fueyo et al., 2021; Gonçalves & Carvalho, 2019; Watson & Matsuzawa, 2018). Since dead infants cannot hold onto their mother’s fur anymore, mothers may transport the corpse in their mouth, ventrally supported by one arm (Bersacola et al., 2025; Carter et al., 2020; Watson & Matsuzawa, 2018), slung across the carrier’s back (Bersacola et al., 2025; Biro et al., 2010), or dragged (Bersacola et al., 2025; Carter et al., 2020; Gonçalves & Carvalho, 2019). Carrying a dead infant hinders foraging and locomotion, thereby increasing predation risks for the carrier (Gonçalves & Carvalho, 2019). Carrying is also energetically costly (Altmann & Samuels, 1992).

Several non-mutually exclusive hypotheses have been put forward to explain why mothers may continue investing in a dead infant (reviewed in Gonçalves & Carvalho, 2019; Watson & Matsuzawa, 2018). Some of these hypotheses imply a concept of death or view infant carrying as a form of grief (Cronin et al., 2011; Takeshita et al., 2020). Other authors do not assume a concept of death (‘unawareness hypothesis’). According to this hypothesis, mothers cannot discern if their infants are dead or temporarily unresponsive and continue to treat them as alive because it is ultimately maladaptive to abandon a non-responsive infant (Sugiyama et al., 2009). Further studies suggest that mothers respond to infantile cues (Alley, 1980) or other cues (fur) that elicit caretaking behaviour (Reiderman et al., 2024). Post-mortem behaviours have also been linked to the post-partum hormonal state of the mother (‘post parturient hypothesis’; e.g., Biro et al., 2010; Kaplan, 1973), and are suggested to be modulated by the bond strength between the mother and the infant (‘mother-infant bond hypothesis’; e.g., Fernández-Fueyo et al., 2021; Matsuzawa, 1997).

Another subset of hypotheses concerns variation due to parity and maternal age. The ‘learning-to-mother hypothesis*’* suggests that nulliparous females are likely to interact with deceased infants of other females to gain experience in mothering (Warren & Williamson, 2004; Watson & Matsuzawa, 2018). The ‘parity hypothesis*’*, in contrast, proposes that multiparous mothers will display behaviours towards dead infants for longer (Sharma et al., 2011), potentially because older mothers place greater value on investing in current offspring (Clutton-Brock, 1984). Das and colleagues (2019) analysed 43 cases across 18 primate species and revealed that older mothers carried dead infants for longer. Another study based on 409 published cases of dead infant carrying in 50 primate species (Fernández-Fueyo et al., 2021) showed that dead-infant carrying was more likely when mothers were younger. However, based on a massive sample with >2000 infants, Sugiyama and colleagues (2009) found that young and older mothers’ dead infant-carrying rates and durations did not differ.

Cannibalism of the dead body has also been described for several primate species. On most occasions, it involves a mother eating (parts of) her dead infant (*filial* or *maternal cannibalism*; e.g., vervet monkey, *Chlorocebus pygerythrus*: Botting & van de Waal, 2020; drills, *Mandrillus leucophaeus*: Casetta et al., 2023; Tonkean macaques, *Macaca tonkeana*: De Marco et al., 2018), but it can also be observed in conspecifics other than the mother (*conspecific necrophagy*; e.g., bonobos, *Pan paniscus*: Fowler & Hohmann, 2010; white-faced capuchins, *Cebus imitator*: Kulick et al., 2021; brown capuchin monkeys, *Sapajus apella*: Trapanese et al., 2020). Cannibalism can provide relatively high-quality food (Milton, 2003), but it may also promote pathogen transmission (Rudolf & Antonovics, 2007). Some authors have suggested environmental stress and abnormal attachment as potential causes (Fedurek et al., 2020), while others consider it an aberrant behaviour (Dellatore et al., 2009) or simply part of a species’ behavioural repertoire (Tokuyama et al., 2017).

To add to the database on post-mortem behaviours toward infants and contribute to broader tests of these hypotheses, we here use twelve years of long-term data to report 22 observations of care-taking behaviours and cannibalism towards dead infants (up to 12 months of age) in a wild population of Guinea baboons (*Papio papio*). Given the lack of prior information on this species, our analyses are exploratory, and we had no specific hypotheses or predictions regarding the occurrence of post-mortem behaviour.

## Methods

### Study species and site

We studied wild Guinea baboons living in the Niokolo-Koba National Park, in Senegal, near the facilities of the Centre de Recherche de Primatologie (CRP) Simenti (13°01’34’’N, 13°17’41’’W) (Fischer et al., 2017). Guinea baboons live in nested multi-level societies. ‘Units’ are at the base of the society and are composed of one ‘primary’ male and one to eight females and their immatures (Dal Pesco et al., 2022; Fischer et al., 2017; unpublished data). Several units group together with bachelor males to form a ‘party’, and two or more parties associate to form ‘gangs’ (Patzelt et al., 2014). Importantly, Guinea baboons show female-biased dispersal, and females may transfer at all levels of the society. The community of Guinea baboons in our study area comprises 350-400 individuals (Dal Pesco et al., 2021; Zinner et al., 2021), of which around 160 individuals in our study population are habituated and identified by physical characteristics. Senegal experiences marked seasonality, with a dry season lasting from November to May and a rainy season from June to October.

### Data collection

We collected data between April 2014 and September 2025 on parties 5, 6 (later split into 6I and 6W), 9 (later split into two parties, with party 9B remaining in our study area), and party 13. During the study period, 61 infants were either seen dead or were presumed dead. We included all behaviours towards dead infants up to an age of 1 year. We augmented our observations using photographs and video. We recorded the behavioural observations using the Samsung Galaxy Note II GT-7100 or Gigaset GX290 handhelds, along with electronic forms designed with Pendragon Software Corporation’s version 7.2.21 software (Chicago, IL, USA). During the study period, the following post-mortem behaviours were observed: carry, protection, grooming, dragging, cannibalism, and meat sharing (Table S1). The information recorded consisted of the date of the event, infant ID (i.e., corpse), the ID of the actor of the post-mortem behaviours, type of relationship between corpse and actor ID (i.e., mother versus non-mother), duration of post-mortem behaviours between actor ID and corpse, and type of post-mortem behaviours observed; all other potentially relevant information was recorded *ad libitum*. Furthermore, we determined the infant’s age at death and the actor’s parity (if the actor was female) from our long-term demographic database, which contains daily information on significant life-history and social events, such as births, deaths, health status, transfers, and female reproductive states. In addition, meteorological data were recorded daily, including minimum and maximum temperatures, minimum and maximum relative humidity, and precipitation.

### Determination of the duration of post-mortem behaviours with a corpse

We calculated the duration of post-mortem behaviours with a dead infant based on the number of days that the researchers could observe the corpse and post-mortem behaviours. For a few cases, we were able to record the total duration of the event, as we saw the mother and the infant alive the day before the post-mortem behaviour began, and the mother without the corpse the day after the last observation of the corpse (Table 1). However, the duration of post-mortem behaviours could not always be accurately determined. Sometimes there was a gap between the last day we saw the mother with the infant alive, the last day we saw the corpse, or the first day we saw the mother again without a corpse, or all three. In these cases, we counted the days the researchers had seen the corpse and estimated only the total duration of behaviour (Table 1); the actual duration of behaviours towards the dead body may have been longer. In five of these cases, we had additional information (e.g., comments in our data about the corpse’s condition) that could suggest a longer duration than the days we had observed in the field.

**Table 1.**
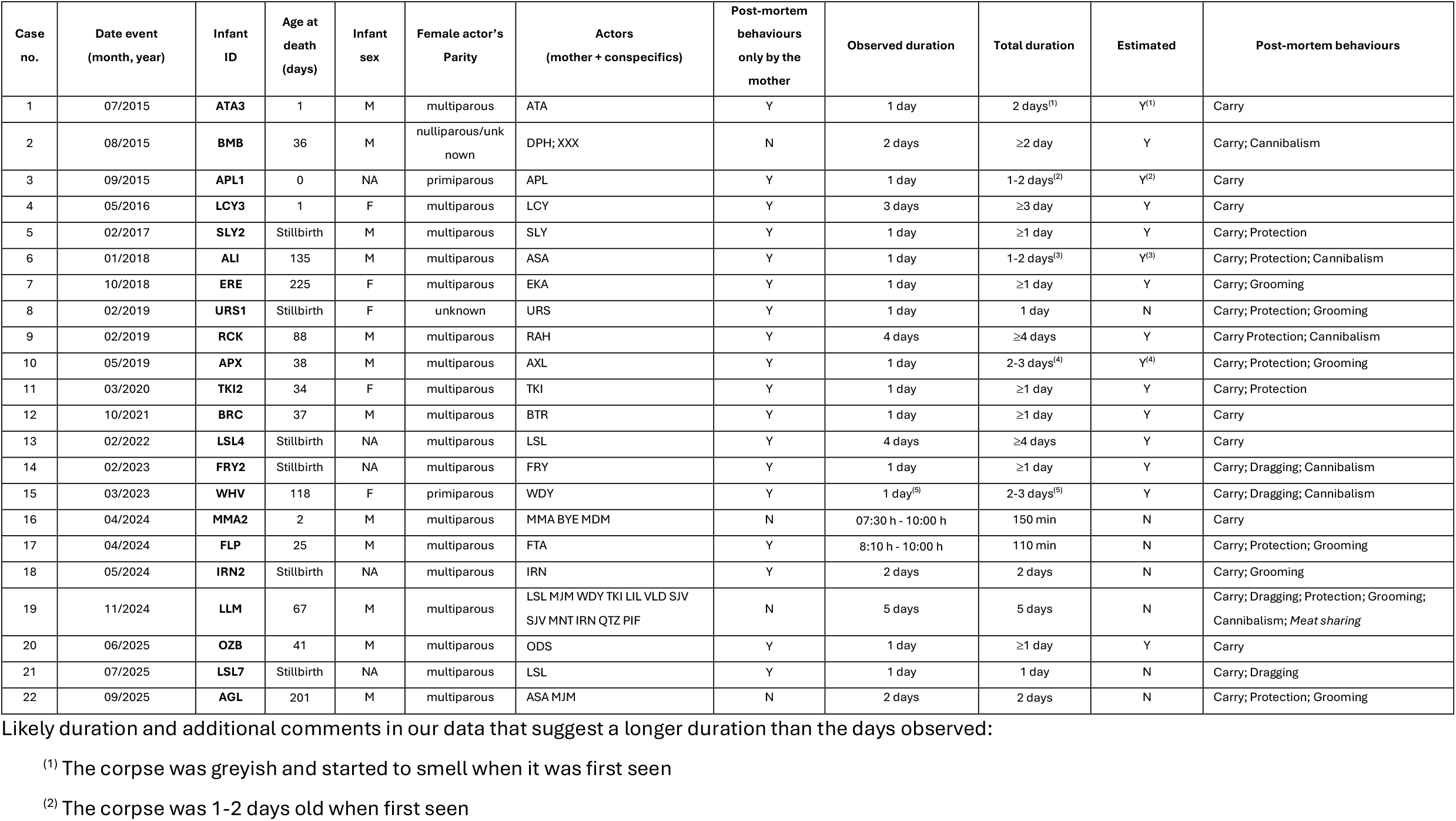
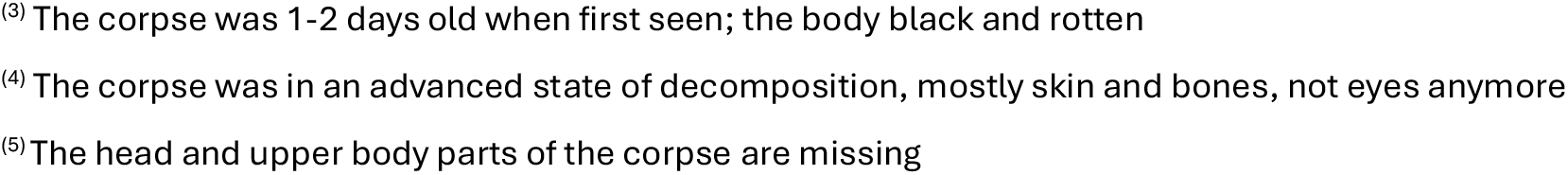
Post-mortem behaviours (N = 22) towards dead infants in a wild population of Guinea baboons from April 2014 to September 2025.

| Case no. | Date event (month, year) | Infant ID | Age at death (days) | Infant sex | Female actor's Parity | Actors (mother + conspecifics) | Post-mortem behaviours only by the mother | Observed duration | Total duration | Estimated | Post-mortem behaviours |
| --- | --- | --- | --- | --- | --- | --- | --- | --- | --- | --- | --- |
| 1 | 07/2015 | ATA3 | 1 | M | multiparous | ATA | Y | 1 day | 2 days<sup>(1)</sup> | Y<sup>(1)</sup> | Carry |
| 2 | 08/2015 | BMB | 36 | M | nulliparous/unknown | DPH; XXX | N | 2 days | ≥2 day | Y | Carry; Cannibalism |
| 3 | 09/2015 | APL1 | 0 | NA | primiparous | APL | Y | 1 day | 1-2 days<sup>(2)</sup> | Y<sup>(2)</sup> | Carry |
| 4 | 05/2016 | LCY3 | 1 | F | multiparous | LCY | Y | 3 days | ≥3 day | Y | Carry |
| 5 | 02/2017 | SLY2 | Stillbirth | M | multiparous | SLY | Y | 1 day | ≥1 day | Y | Carry; Protection |
| 6 | 01/2018 | ALI | 135 | M | multiparous | ASA | Y | 1 day | 1-2 days<sup>(3)</sup> | Y<sup>(3)</sup> | Carry; Protection; Cannibalism |
| 7 | 10/2018 | ERE | 225 | F | multiparous | EKA | Y | 1 day | ≥1 day | Y | Carry; Grooming |
| 8 | 02/2019 | URS1 | Stillbirth | F | unknown | URS | Y | 1 day | 1 day | N | Carry; Protection; Grooming |
| 9 | 02/2019 | RCK | 88 | M | multiparous | RAH | Y | 4 days | ≥4 days | Y | Carry Protection; Cannibalism |
| 10 | 05/2019 | APX | 38 | M | multiparous | AXL | Y | 1 day | 2-3 days<sup>(4)</sup> | Y<sup>(4)</sup> | Carry; Protection; Grooming |
| 11 | 03/2020 | TKI2 | 34 | F | multiparous | TKI | Y | 1 day | ≥1 day | Y | Carry; Protection |
| 12 | 10/2021 | BRC | 37 | M | multiparous | BTR | Y | 1 day | ≥1 day | Y | Carry |
| 13 | 02/2022 | LSL4 | Stillbirth | NA | multiparous | LSL | Y | 4 days | ≥4 days | Y | Carry |
| 14 | 02/2023 | FRY2 | Stillbirth | NA | multiparous | FRY | Y | 1 day | ≥1 day | Y | Carry; Dragging; Cannibalism |
| 15 | 03/2023 | WHV | 118 | F | primiparous | WDY | Y | 1 day<sup>(5)</sup> | 2-3 days<sup>(5)</sup> | Y | Carry; Dragging; Cannibalism |
| 16 | 04/2024 | MMA2 | 2 | M | multiparous | MMA BYE MDM | N | 07:30 h - 10:00 h | 150 min | N | Carry |
| 17 | 04/2024 | FLP | 25 | M | multiparous | FTA | Y | 8:10 h - 10:00 h | 110 min | N | Carry; Protection; Grooming |
| 18 | 05/2024 | IRN2 | Stillbirth | NA | multiparous | IRN | Y | 2 days | 2 days | N | Carry; Grooming |
| 19 | 11/2024 | LLM | 67 | M | multiparous | LSL MJM WDY TKI LIL VLD SJV<br>SJV MNT IRN QTZ PIF | N | 5 days | 5 days | N | Carry; Dragging; Protection; Grooming; Cannibalism; *Meat sharing* |
| 20 | 06/2025 | OZB | 41 | M | multiparous | ODS | Y | 1 day | ≥1 day | Y | Carry |
| 21 | 07/2025 | LSL7 | Stillbirth | NA | multiparous | LSL | Y | 1 day | 1 day | N | Carry; Dragging |
| 22 | 09/2025 | AGL | 201 | M | multiparous | ASA MJM | N | 2 days | 2 days | N | Carry; Protection; Grooming |

2 Likely duration and additional comments in our data that suggest a longer duration than the days observed:
<sup>(1)</sup> The corpse was greyish and started to smell when it was first seen
<sup>(2)</sup> The corpse was 1-2 days old when first seen
<sup>(3)</sup> The corpse was 1-2 days old when first seen; the body black and rotten <sup>(4)</sup> The corpse was in an advanced state of decomposition, mostly skin and bones, not eyes anymore <sup>(5)</sup> The head and upper body parts of the corpse are missing

### Compliance with ethical standards

All applicable international, national, and/or institutional guidelines for the care and use of animals were followed. Research was conducted in accordance with the regulations set by the Senegalese agencies and the ethical standards of the Animal Care Committee at the German Primate Centre. This study followed the ASAB guidelines for the ethical treatment of nonhuman animals in behavioural research (ASAB Ethical Committee/ABS Animal Care Committee, 2023). Approval and research permission were granted by the DPN and the MEPN de la République du Sénégal.

## Results

During the study period, 67 infants died before reaching 1 year of age. In 4 cases, we could not establish when the infants had died because the field station was closed due to the COVID-19 pandemic. In 22 of the remaining 63 cases, mothers, but occasionally also other group members, carried, protected, groomed, and dragged dead infants (summarised in Table 1). The distribution of these events was not restricted to a particular season, with 8 cases (36%) observed during the rainy season and 14 cases (64%) during the dry season (Table 1), which roughly corresponds to the relative lengths of the rainy (42%) and dry seasons (58%). Most cases (20 out of 22) involved infants up to 6 months old (Table 1). Of these infants, six were stillbirths, four were less than a week old, eight were between one week and three months old, and two were between three and six months old. The two remaining cases were infants older than 6 months; therefore, they had already transitioned to “brown” infants (Table A1), with the oldest infant dying at 225 days. The average age at death of infants for whom we observed post-mortem infant-directed behaviours was 46.4 days (N = 22), whereas infants for whom we did not observe such behaviours were on average 114.7 days old (N = 41).

The behaviours towards the corpse lasted from a few hours to five days (Table 1). The behaviours observed were dead infant carrying (Fig. 1), protection, grooming (Fig. 2), dragging (Fig. S3.1C), and cannibalism (Fig. 3). Carrying occurred in all 22 cases. The corpse was carried in the actor’s arm, either ventrally supporting it (Fig. 1A), holding it in one hand (Fig. 1B), or in the mouth (Fig. 1C). When dragging occurred, the head of the dead infant was frequently banged against stones or rocks. Cannibalism was seen on six occasions (27.3%). It consisted of eating the gums and tongue of the infants, sometimes pulling the meat with the teeth to extract it (Fig. 3A), except in case 19, in which the mother was also seen eating other parts (Fig. 3B), such as the brain of the infant after opening the skull, and the meat of the mandible (see supplement section S3 for details in the chronology of events of case 19). Neither dragging nor cannibalism was seen when the infant had recently died; instead, these behaviours occurred when the body had started to decompose. In the few cases in which dragging and cannibalism occurred the first day the corpses were observed, the advanced state of the infant’s decomposition and the absence of observations in the previous days suggest that we had missed the beginning of the event.

**Figure 1.**
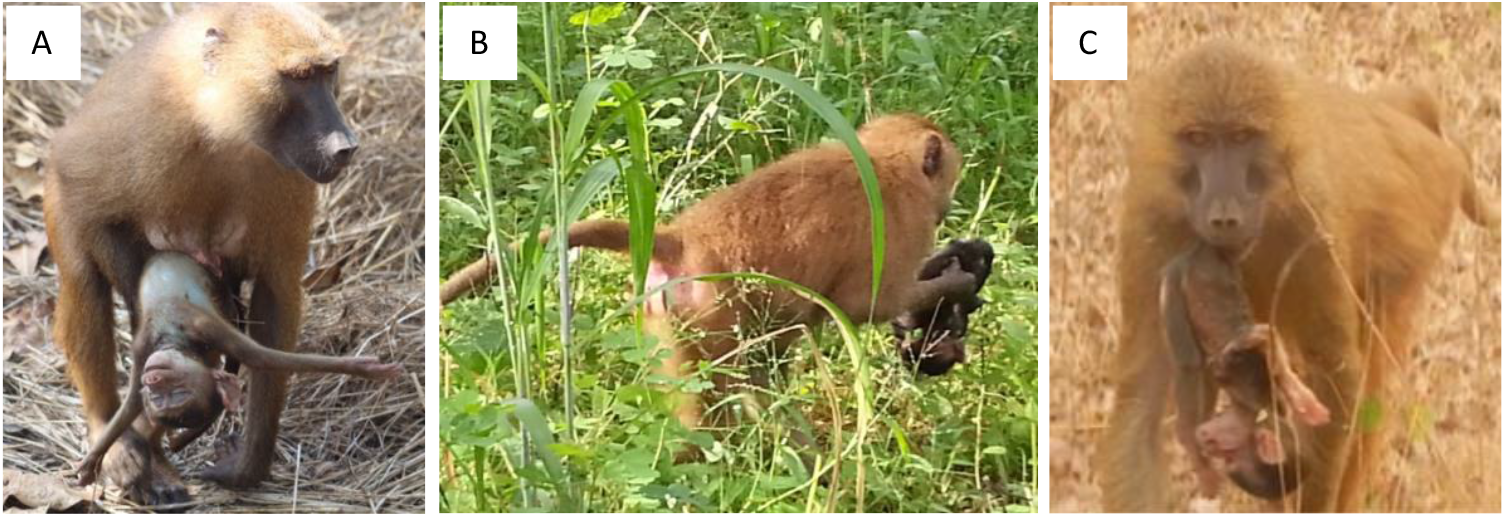
The modes of carrying a dead infant in Guinea baboons include (A) ventrally supporting the corpse with one arm (case 19: LLM), (B) holding it in one hand (case 1: ATA3), and (C) in the mouth (case 5: SLY2). Picture A taken by Anaïs Avilés de Diego; 11/2024, picture B taken by Federica Dal Pesco, 09/2015, and picture C taken by Lauriane Faraut, 02/2017; Cognitive Ethology Laboratory, German Primate Centre.

**Figure 2.**
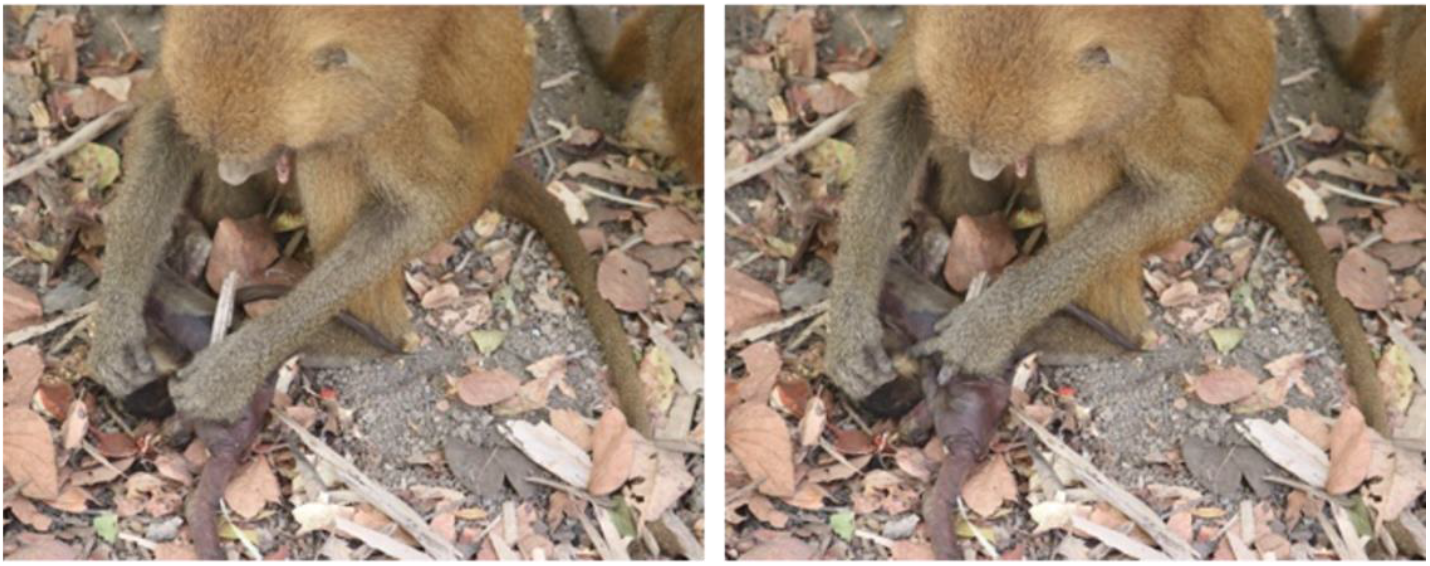
Dead infant APX being grooming by his mother (case 10). Pictures taken by Anaïs Avilés de Diego, 05/2019; Cognitive Ethology Laboratory, German Primate Centre.

**Figure 3.**
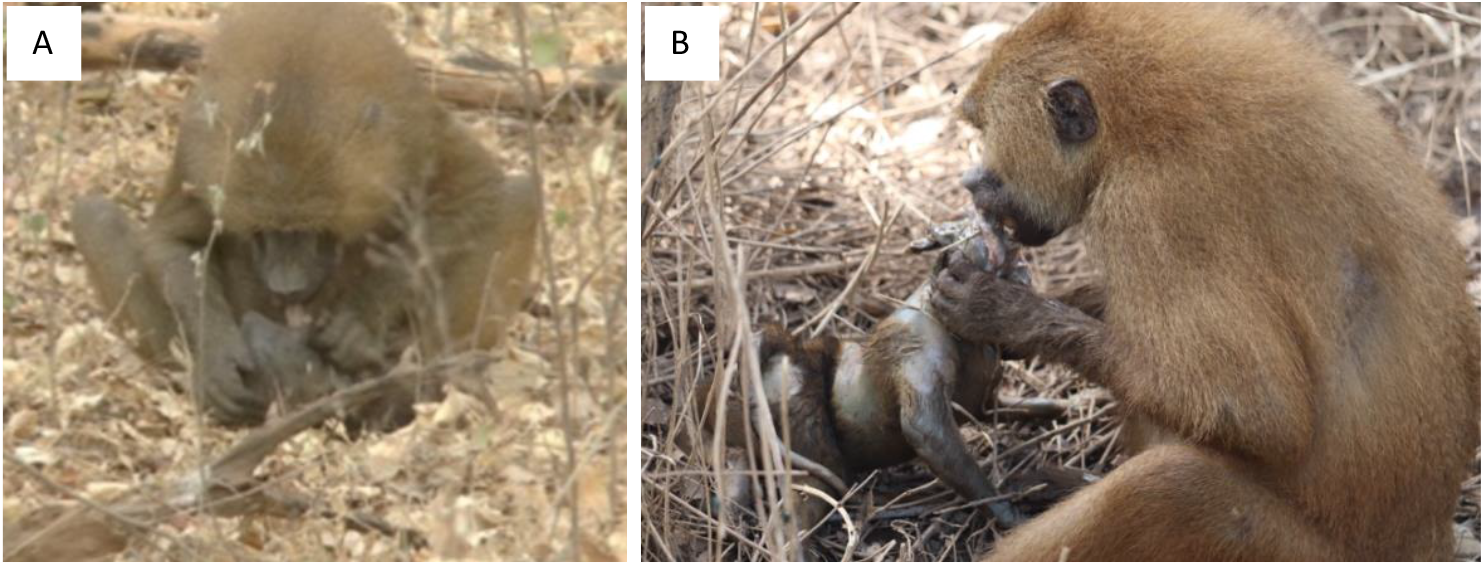
Dead infants being cannibalised by their mothers. In picture (A) the mother is consuming the tongue of the infant corpse (case 6: ALI), whereas in picture (B) the mother is eating parts of the dead infant’s brain (case 19: LLM). Picture A is from 01/2018 and B from 11/2024, both taken by Anaïs Avilés de Diego; Cognitive Ethology Laboratory, German Primate Centre.

In 18 out of 22 cases, the only actors of the post-mortem behaviours were the mothers of the dead infants. In the remaining three instances (cases 16, 19, and 22), other individuals besides the mother also showed post-mortem behaviours with the corpses, being mostly members from the same unit and party of the dead infant, but also by other individuals who belonged to different party but had at previously shared unit or party with the dead infant’s mother (case 19). In case 16, the older brother of the dead infant touched and carried the corpse, and the primary male of the mother (and potential father of the infant) handled the infant and grunted at it. Case 19 was the longest observed, with many individuals displaying behaviours towards the corpse besides the mother, for instance, the dead infant’s older sisters, and another female (from another party) who also cannibalised the infant (Fig. S3.4). None of the actors involved showed any visible signs of distress, such as contact call vocalisations.

Eighteen of the 22 dead infants had multiparous mothers; two dead infants had primiparous mothers, and for one infant, maternal parity was unknown. In the remaining case (case 2), the mother had died, and the female who carried the dead infant was nulliparous, while the second female, who cannibalized it, was of unknown parity. The cause of death was unknown for all infants, but we suspected a snake bite in case 19 (Fig. S2.1), and an injury after the baboons had passed through a particularly thorny area in case 11 (Fig. S2.2). Infanticide seems unlikely since it has never been observed in our population of Guinea baboons, and pregnant and lactating females may transfer to a new primary male without any known following feticide or infanticide occurrences.

## Discussion

Over twelve years of observation (2014-2025), we observed post-mortem behaviours directed towards 22 dead infants, which amounts to around a third of the 63 total cases where we could record the age of death. For infants for whom we did not have records of infant-directed post-mortem behaviours, we cannot determine whether we may have missed such behaviours because we did not follow the group on that particular day or the next, whether the mothers simply abandoned the corpse, or whether the infants were preyed upon by birds, snakes, or other predators. As the average age of infants for whom we observed no post-mortem behaviours was substantially higher (114.7 days) than that of those for whom we did (46.4 days), predation may have indeed played a role. With increasing age, infants gradually spend more time away from their mothers (Avilés de Diego et al., 2026) and face an increasing risk of predation, although we have never directly observed any attacks.

Dead infant carrying was the predominant post-mortem behaviour, consistent with previous reports in primates, followed by protection, grooming, and dragging. Cannibalism also occurred. Since Guinea baboons opportunistically hunt and feed on prey (Goffe & Fischer, 2016; O’Hearn et al., 2025), meat consumption is not unusual. The duration of behaviours towards dead infants in this study (<1 to 5 days) is similar to the one of reported in other baboon species (e.g., yellow baboons, *Papio cynocephalus*, ca. 3 days: Altmann, 1980; chacma baboons, 2-10 days: Carter et al., 2020; Cheney & Seyfarth, 2007), and short compared to the maximum registered in other primate species, (e.g., Tonkean macaques, 25 days: De Marco et al., 2018; geladas, *Theropitecus gelada*, 48 days: Fashing et al., 2011; Hanuman langurs, *Semnopithecus entellus*, 27 days: Sharma et al., 2011; chimpanzees, 89 days: Soldati et al., 2022; and four months: Hosaka et al., 2000). Carter and colleagues proposed that the cost of carrying a dead infant in species that travel long daily distances could lead to shorter dead-infant carrying durations (Carter et al., 2020). The long daily travel distances of the Guinea baboons (on average 3.5 km, up to 12 km (Zinner et al., 2021) may promote a relatively early abandonment of dead infants.

The dead infants were carried in the actor’s arm, either ventrally supporting them, holding them in one hand, or, less frequently, in the mouth. Guinea baboon mothers do not carry live infants in one hand or in the mouth, nor do they drag them on the floor. However, food items such as fruit or prey can be held in the hand or mouth, or even dragged while moving or travelling. The high occurrence of care-taking behaviours at the onset of the interactions with a corpse and the eventual decline of these behaviours, along with the sequential changes in the way of carrying – from a type typical from an alive infant to a type seen with food (Fig. S3.1) – suggest that the infant corpse might initially be perceived as something that requires care-taking behaviours but progressively loses this characteristic, being finally perceived as a food item.

Given that most of the mothers that showed post-mortem behaviours were multiparous, this finding speaks against the idea that dead infant carrying is mainly driven by the need to learn to mother (Warren & Williamson, 2004; Watson & Matsuzawa, 2018). There was also no evidence in support of the mother-infant bond hypothesis (Fernández-Fueyo et al., 2021; Matsuzawa, 1997), because mothers were not more likely to carry the corpse of older infants than newborns. A recent study on Mountain gorilla responses found no effect of parity or season on “infant corpse carrying” (ICC) (Munyawera et al., 2026). However, among non-maternal carriers, nulliparous and primiparous females were more likely to show ICC than multiparous females, which the authors interpreted as evidence for the ‘learning-to-mother’ hypothesis (Munyawera et al., 2026).

We suggest that the most parsimonious explanation for the Guinea baboons is a combination of the infantile cue and the unawareness-of-death hypothesis. The lack of infant responses to maternal behaviour and the disintegration of the corpse may drive the transition from cues that evoke caretaking to cues associated with food, which ultimately facilitates occurrences of cannibalism. Watson and Matsuzawa (2018) also suggested that cannibalism – once the corpse had started to disintegrate – could indicate a weakening of the mother-infant bond and a transition from regarding the corpse as alive to an object/food item, and that the overlap between caring and cannibalism that occasionally occurs might indicate conflicting impulses (Watson & Matsuzawa, 2018). In support of this view, we have also observed grooming of prey after a hunting event, suggesting that fur per se may elicit grooming activity (‘mammalian cues hypothesis’: Reiderman et al., 2024).

Taken together, the Guinea baboons’ behaviour towards dead infants is best explained as cue-driven, rather than concept-driven. In other words, the observed behaviour does not require an understanding of the irrevocability of death. Our study adds to the growing body of evidence on animals’ responses to dead conspecifics and contributes to comparative analyses of primate thanatology (Arlet et al., 2025).

## Supporting information

ESM_GB_post-mortem

## Acknowledgements

We are grateful to the Diréction des Parcs Nationaux (DPN) and the Ministère de l’Environnment et de la Protéction de la Nature (MEPN) de la République du Sénégal for approval to conduct this study in the Parc National du Niokolo-Koba (PNNK). We particularly thank the all the conservators of the park during the study period (Oussoumane Kane, Mallé Gueye, Amar Fall, Assane Ndoye, Jacques Gomis, Paul Moïse Diedhiou, and Ibrahima Ndao) and deputy conservator Elhadji Mamadou Thiaw for their cooperation and support. We thank all the CRP Simenti field assistants and students, in particular Cheikh Sané, Moustapha Dieng, Moustapha Faye, Armel Louis Nyafouna, El’Hadji Yankhoba Dansokho, Touradou Sonko, Vieux Biaye, Djibril Coly, Amadou Bamba Diedhiou, Chérif Younousse Kéba Camara, Jean Louis Diouf, Lassana Ba, Jean Malack, Boubacar Sow, Malamine Diedhiou, Moussa Dieng, Ndiouga Diakhate, Dame Diallo, Hamidou Sogoba, and Alfred Oga, for their support and work in the field. This research was supported by the Deutsche Forschungsgemeinschaft (DFG, German Research Foundation), Research Training Group 2070 “Understanding social relationships”, Grant/Award Number: 254142454 /GRK 2070.

## Author contributions

Conceptualization: AAdD, FDP, JF; Investigation: AAdD, FDP, and the CRP Simenti field team; Methodology: AAdD, FDP, JF; Data curation: AAdD, FDP, JF; Writing –Original Draft: AAdD; Writing – Review & Editing: AAdD, FDP, JF; Visualization: AAdD, FDP, JF; Funding acquisition: JF.

## Data availability statement

All data generated or analysed during this study are included in this published article.

