## Supplementary material for "Post-mortem infant-directed behaviours in wild Guinea baboons": ESM_GB_post-mortem

### ^1^Department for Primate Cognition, Georg-August-Universität, Göttingen

### ^2^Cognitive Ethology Laboratory, German Primate Centre, Göttingen

### *Author responsible for correspondence: Anaïs Avilés de Diego

### Address: Cognitive Ethology Laboratory, German Primate Centre, GmbH, Kellnerweg 4, 37077 Göttingen, Germany.

### Telephone number: +49 551 3851-248

### S1. Ethogram

**Table S1**. Ethogram of post-mortem behaviours that were directed to Guinea baboon infant corpses from April 2014 to September 2025.

| **Behaviour** | **Definition** |
| --- | --- |
| Carry | The infant corpse is carried by its mother (or another individual), ventrally supported with one arm, held in one hand, and in the mouth |
| Protection | The mother (or another individual) holds or grabs tightly the infant corpse with one or both arms and/or feet, potentially intensifying when other members are in proximity, thereby limiting the access of other individuals to the corpse or pulling back if other members try to get hold of it. |
| Grooming | Movement of fingers and/or lips through the fur and the skin of the infant corpse by the mother (or another individual). |
| Dragging | The mother (or another individual) pulls the infant corpse across the ground by grabbing it by the one of its extremities, tail, or torso. |
| Cannibalism | The mother (or another individual) consumes the meat of the infant corpse. |
| Meat sharing | The mother (or another individual), who is consuming the dead infant’s meat, tolerates the removal of pieces of meat from the dead infant meat items from the dead infant from the possessor (*passive meat sharing*; see Goffe & Fischer, 2016). |

See the CRP Simenti work manual (Dal Pesco & Fischer, 2022) for age category definitions and other behaviours mentioned in the text.

### S2. Infant injuries

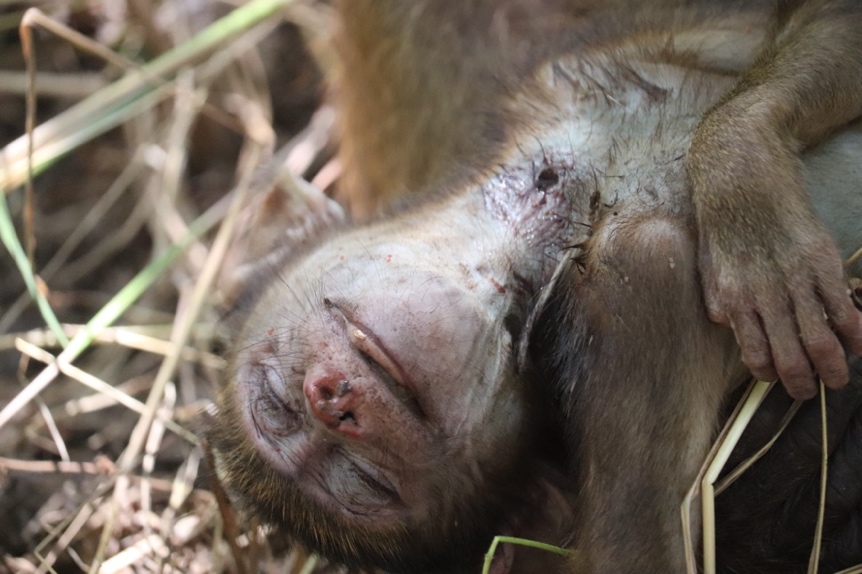

**Figure S2.1.** Wound at the base of the neck of the dead infant LLM, which might have been caused by a snake bite (see the two rounded spots with necrotic tissue around). Picture taken by Anaïs Avilés de Diego on 04/11/2022; Cognitive Ethology Laboratory, German Primate Centre.

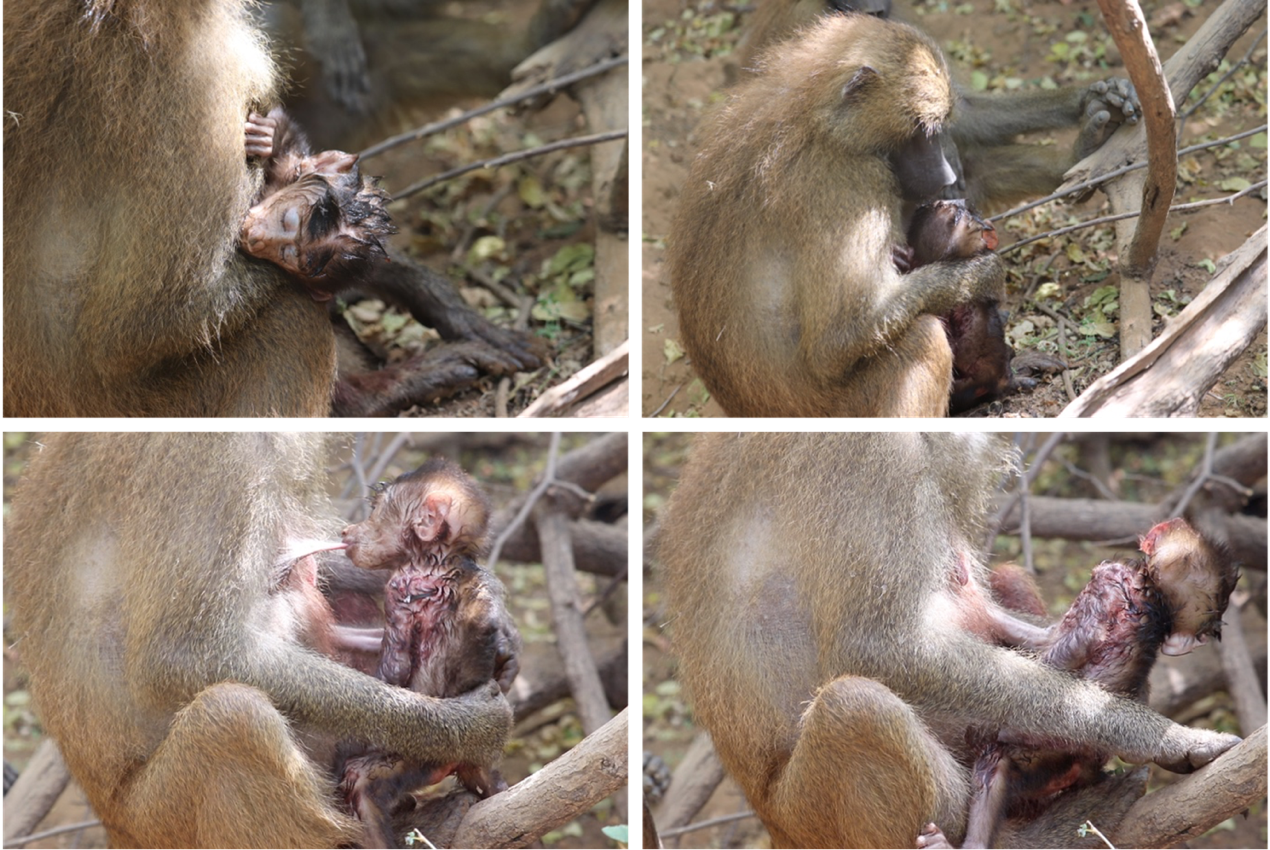

**Figure S2.2.** Injured infant TKI2 with its mother (case 11) after the baboons had passed through a particularly thorny area. Blood is visible on the left side of the infant’s body. Pictures taken by Anaïs Avilés de Diego on 03/2020; Cognitive Ethology Laboratory, German Primate Centre.

### S3. Case 19: Chronology of events LSL-LLM

Case 19 in particular gave us a partial glimpse into the potential time sequence of post-mortem behaviours with a corpse and potential changes in mental representation: first, the way of carrying changed throughout the event, from being ventrally supported by the mother, to being carried with one hand, in the mouth, and finally being dragged on the floor, all from a careful to a careless manner (Fig. S3.1). Care-taking also decreased, with grooming and protection higher on the first day, then reducing throughout the second day, and not occurring on the remaining days. In case 19 we could see a clear switch in the mother behaviour, who was carefully interacting with the dead infant until she started cannibalized it, starting with the gums and tongue, and then opening the skull and eating the brain, and meat-sharing with another female in a similar way to when Guinea baboons share the meat of a prey after an opportunistic hunting event. The mother continued carrying the corpse in a more or less careful manner, the day that she had started cannibalising, and still groomed him afterwards, which might indicate this temporary double mental representation of the corpse as infant and as meat.

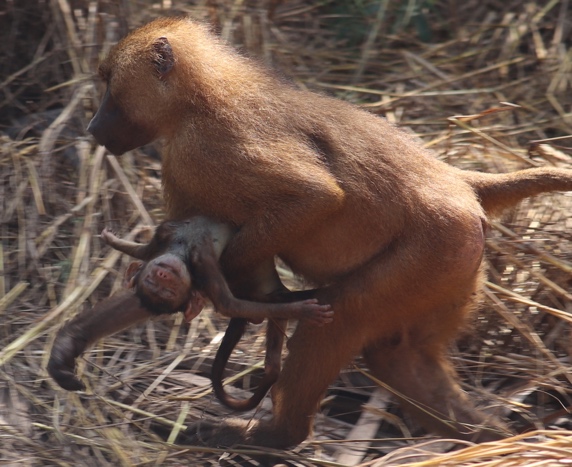

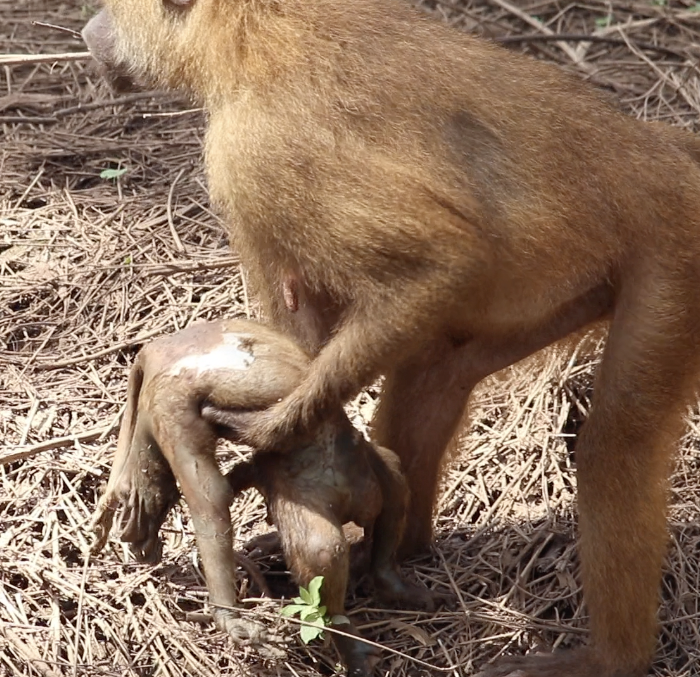

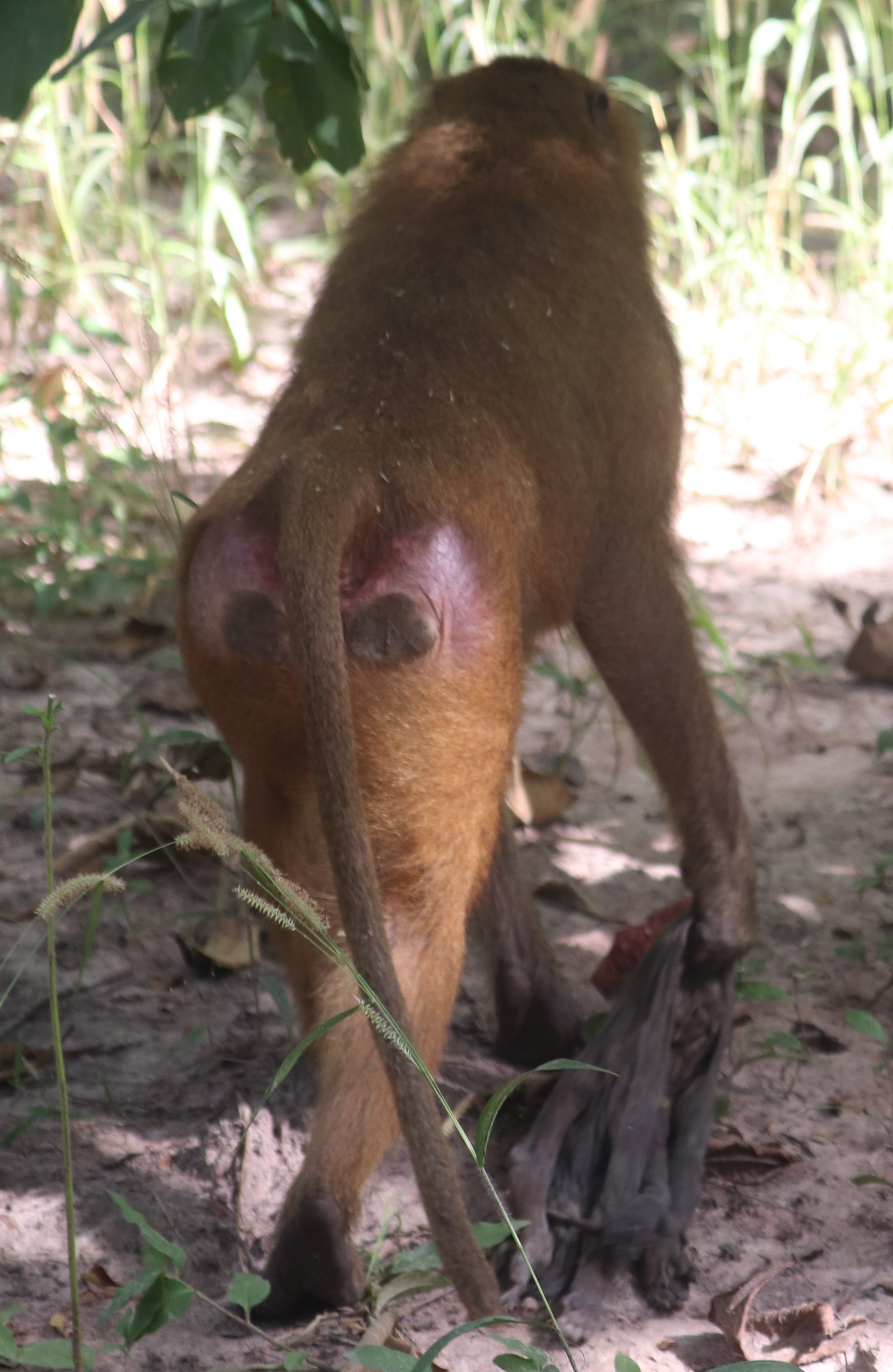

A

B

C

**Figure S3.1.** LLM is initially transported by his mother more carefully, but the carrying changes over the following days to a more careless manner. On day 1 (A), the mother supported the corpse ventrally, on day 2 (B), she held it with one hand, and on day 3 (C), she dragged it on the floor. See the lack of hair in some areas of the body in picture (B), and the lack of hair and body mass in picture (C). Pictures taken by Anaïs Avilés de Diego on 04-05-06/11/2024; Cognitive Ethology Laboratory, German Primate Centre.

**Day 1**

On 04/11/2024, the researchers found the baboon party at 06:47 h. The mother (LSL) was found with her dead black infant male (LLM). LLM still had the black natal coat and light pinkish skin coloration. The death of LLM seemed very recent (a few hours earlier), since the corpse looked very fresh: the skin was still pinkish and did not lack hair; there were no signs of putrefaction: no disfiguration of the face or body, no bloating, nor a foul smell. Moreover, LLM had a wound in the upper part of the back of the head that looked very recent: the blood was bright red and slightly runny; however, it is unclear if the wound was from before death, done post-mortem, or if it was the cause of death. Two black, rounded spots were seen in the base of the neck of the infant (Fig. S2.1), and the skin around them was necrotic, which could hint at a snake bite as a possible cause of death. We established the date of death on 04/11/2024, presuming that the infant had died during that same morning. Throughout the day, LSL maintained contact with or kept LLM within close proximity (<1m), and carried LLM carefully, ventrally supporting the corpse with one hand (Fig. S3.1A). LSL also groomed and cradled LLM. LSL started swatting away flies from LLM at some point in the morning, and this continued as the day went on. LSL was protective of the corpse sometimes, mostly of her primary male VLD (subadult male) and SNV (subadult female). The latter grabbed LLM several times and tried to hold him, but LSL firmly held LLM by the arm and leg and pulled him back. SNV groomed LLM on one occasion when LSL was being protective and impeding SNV from getting LLM. Other individuals also interacted with LLM. For instance, the older sisters of LLM approached (two juvenile females), and an infant handled the corpse, and the youngest one also touched, groomed, embraced, and inspected LLM.

**Day 2**

On 05/11/2024, the baboon party was found at 06:56 h. The dead infant was still with LSL. The skin of LLM was greyish, darker in the face, and several parts of the body presented absence of hair: most of the head and face, some areas in the chest, abdomen, arms, inner legs, and some patches of the tail (Fig. S3.1B). When LSL was carrying LLM, his head and spine were bending back and forth, further compared to the previous day. The body of LLM appeared bloated, and the penis (Fig. S3.2) and the belly button were exceedingly swollen. LLM had a pungent smell (this was the day that the corpse smelled more intensely), and there were many flies around it. The skull bones seemed cracked under the skin (Fig. S3.2), and the upper part of the back of the head had a white-reddish mark, which was most likely the area of the wound of the day before. The two black, rounded spots at the base of the neck did not appear significantly different from the day before; if anything, they were less perceivable.

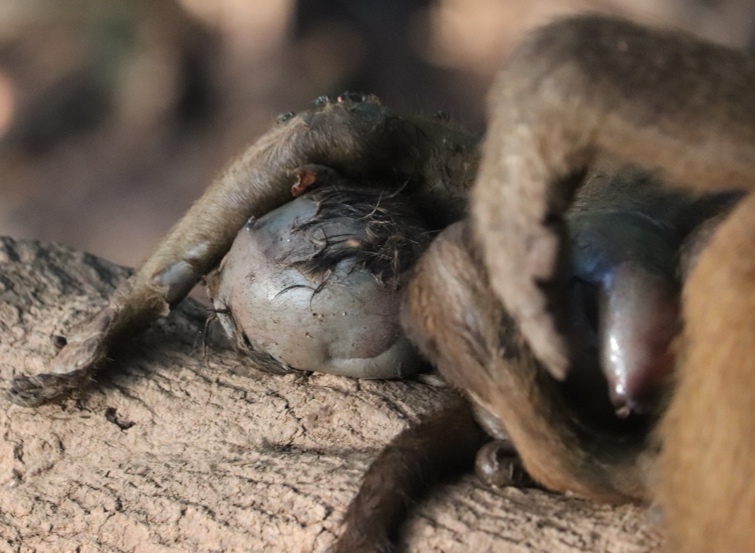

**Figure S3.2.** Dead infant LLM at the beginning of day 2. The corpse had a white-reddish mark on the upper part of the back of the head, and the skull bones appeared cracked underneath. The swollen penis is visible in the right area of the picture. Picture taken by Anaïs Avilés de Diego on 05/11/2024; Cognitive Ethology Laboratory, German Primate Centre.

In the early morning, LSL carried LLM ventrally, supporting the corpse with one hand. LSL seemed to carry the LLM less in contact with her torso and less carefully than the day before. LSL groomed LLM occasionally, but much less frequently than the previous day (grooming occurred mostly in the early morning on day 2). Whenever LSL was sitting, she had LLM at her feet and frequently swatted flies away. At around 09:00h, the number of flies had considerably increased, and so did the frequency with which LSL swatted flies away. During the early morning, QTZ, a mid-juvenile male, approached, sniffed both LLM and LSL’s mouth, grunted, and shortly groomed LLM’s tail. LSL avoided VLD once when he tried to approach, and she was protective of LLM by strongly holding it.

At around 10:10, LSL was first seen cannibalising LLM: she first ate from the gums and tongue. Shortly after, LSL sat in an area with dense vegetation, almost out of sight of the other baboons and the researchers. In this moment, the cranial cavity of LLM was exposed, and LSL was eating parts of LLM’s brain. Once LSL came out of the dense area, she continued eating from LLM’s brain. At around 10:20, LSL was holding a loose piece of skull and eating parts of LLM’s brain adhered to it (Fig. S3.3). Then, LSL dropped the loose piece and directly ate what was left of the brain inside LLM’s skull, while grabbing LLM by the head, and filling up her cheek pouches.

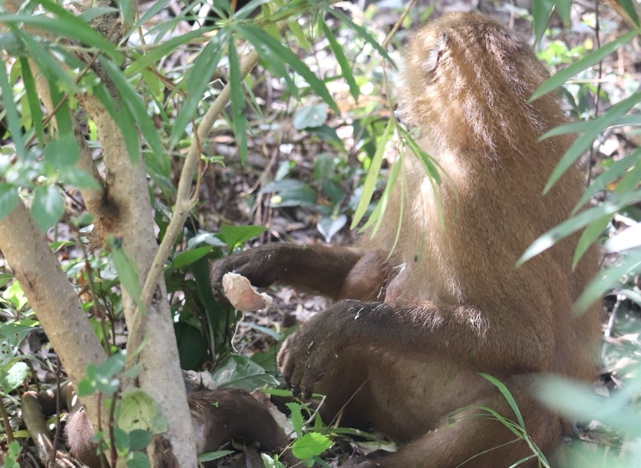

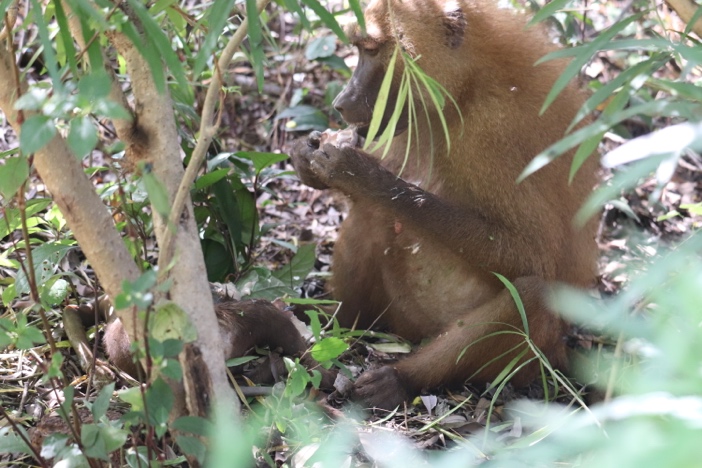

**Figure S3.3.** Dead infant LLM being cannibalised by his mother during the morning of day 2. The mother has a piece of skull in her hands, and she is eating the brain matter attached to it. Picture taken by Anaïs Avilés de Diego on 05/11/2024; Cognitive Ethology Laboratory, German Primate Centre.

At 10:21 h, the group started moving. LSL carried LLM, and whenever she stopped walking, she continued cannibalising LLM. The transport of LLM was now more careless compared to earlier that day or to the day before, and she held LLM with one hand but did not ventrally support it (Fig. S3.1B), similar to when Guinea baboons carry food or prey. At 10:50 h, LSL carried LLM in her mouth for the first time, but she then returned to carrying LLM in her hands. LSL was also observed dropping LLM carelessly on the ground. Eventually, it was possible to see that the whole LLM’s upper cranial vault and its content were missing, with just some skin left. Throughout the day, LSL continued swatting away flies occasionally.

Once LSL started eating LLM’s brain, some individuals approached her and sniffed and/or looked in her direction. VLD approached LSL-LLM within two meters, and some seconds later, LSL moved several meters away from VLD while guarding LLM. LSL continued eating once she sat down again.

At 10:50 h, LSL and IRN (mature adult female) were sitting next to each other at <1m of proximity, and LSL was eating two loose pieces of LLM’s skull and having LLM’s body next to her. LSL then dropped both these pieces on the ground. Immediately, while LSL was still within 1m, IRN took one of these pieces of bone (Fig. S3.4). IRN was in full view of LSL when this occurred, being this the first occurrence observed in Guinea baboons of a mother sharing the meat of her deceased infant with another individual.

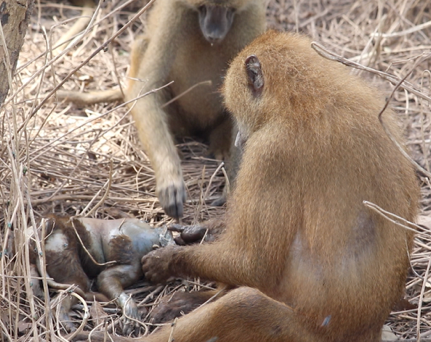

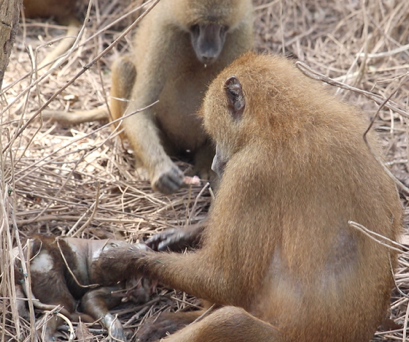

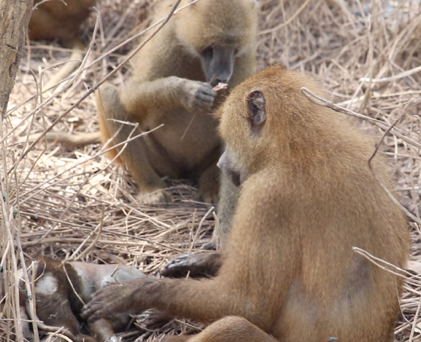

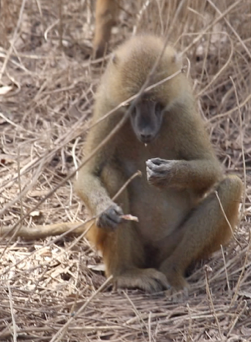

**Figure S3.4.** Progression of a meat sharing event with the meat of a dead infant. The mature adult female IRN is at the back, and she is grabbing and manipulating a piece of meat from the dead infant LLM that its mother LSL dropped on the floor. Picture taken by Anaïs Avilés de Diego on 05/11/2024; Cognitive Ethology Laboratory, German Primate Centre.

For the rest of the day, LSL continued carrying LLM whenever the group moved by holding it in one hand, and sitting down from time to time and dropping LLM at her feet, either in contact or in close proximity (< 1m), eating from her cheek pouches or picking up some pieces of meat from LLM’s head and eating them, and swatting flies away. Eventually, LSL broke LLM’s mandible and ate the meat attached to it. Other individuals remained in the vicinity of the dyad, occasionally approaching and sniffing LSL’s mouth. At 12:02, LSL was seen eating a coconut, this being the first time after cannibalism had started that she was eating a food item that was not LLM. From then on, LSL continued eating pieces of LLM, but less frequently than before. The researchers left the baboon party at 12:45 h. By then, LLM had almost no hair on its torso and arms; it was still slightly bloated and smelled strongly.

**Day 3**

The researchers found the baboon party on 06/11/2024 at 08:14 h in an area with very dense vegetation. LSL still had LLM, but she was relatively separated from the main group, about 15m behind. The skin of LLM was dark grey, partially torn, and hairless (Fig. S3.1C). The head was totally unrecognisable, and the bones of the torso seemed to be missing (Fig. S3.1C). LLM had lost most resemblance to an infant. The body of LLM was no longer bloated and had a foul smell, though not as strong as the day before. There were no apparent flies around, but the dense area might have made it difficult to see.

The baboons spent their morning foraging in the dense area. LSL was observed a couple of times with LLM in her hand, but she mostly dragged LLM along the ground (Fig. S3.1C). LSL picked parts of LLM’s belly with her fingers and ate them, but cannibalism occurred less frequently than the day before. On occasions, LSL sat directly on top of LLM. None of the group members showed any behaviour towards the dead infant; LSL was rather away from the group, and no one approached or seemed to have any interest in her and the dead infant (only a mid-juvenile sniffed the corpse). On one occasion, when VLD was passing by, LSL avoided him, but no protective behaviour was observed. The researchers left the baboon party at 12:14 h.

**Day 4**

The researchers did not find the baboon party to which LSL-LLM belonged on 07/11/2024.

**Day 5**

When the researchers found the baboon party again at 06:52 h on 08/11/2024, LSL was still holding on to LLM, who was barely a piece of dry, leathery skin (Fig. S3.5). There were no head or visible body parts anymore, and it had a foul smell. Cannibalism occurred, but with very low frequency, and since the corpse did not look like an infant anymore, it was very difficult to determine from which part LSL ate. Throughout the day, LSL held LLM firmly, not letting go, for instance, when grooming another individual (Fig. S3.5). On this day, LSL was not seen carrying LLM; she was only dragging him along the ground. No protective behaviour was observed from LSL, and no individual showed any post-mortem behaviour towards LLM. The researchers left the baboon party at 13:02 h.

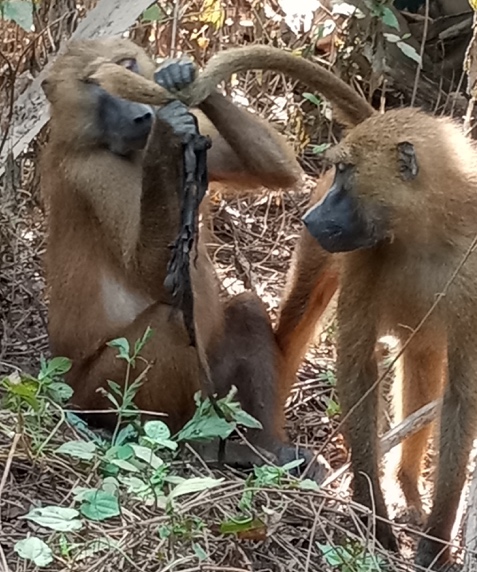

**Figure S3.5.** LSL holding the dead infant LLM while grooming another individual. Picture taken by Eva Rouselle on 08/11/2024; Cognitive Ethology Laboratory, German Primate Centre.

**First day without seeing the corpse**

On 09/11/2024 the researchers found the baboon party at 07:02h, and on this day LSL did not have the dead infant LLM with her anymore.
